# Interactions between supervised and reinforcement learning processes in a neurorobotic model

**DOI:** 10.1101/2022.09.30.510289

**Authors:** Adriano Capirchio, Chiara Ponte, Gianluca Baldassarre, Francesco Mannella, Elisa Pelosin, Daniele Caligiore

**Affiliations:** AI2Life s.r.l., innovative start-up, Institute of Cognitive Sciences and Technologies, National Research Council spin-off, Via Sebino 32, Rome, 00199, Italy; Computational and Translational Neuroscience Laboratory, Institute of Cognitive Sciences and Technologies, National Research Council, Rome, Italy; Department of Neuroscience, Rehabilitation, Ophthalmology, Genetics, Maternal and Child Health, University of Genoa, Genoa, Italy; Laboratory of Embodied Natural and Artificial Intelligence, Institute of Cognitive Sciences and Technologies, National Research Council, Rome, Italy; Institute of Cognitive Sciences and Technologies, National Research Council, Rome, Italy; Ospedale Policlinico San Martino, IRCCS, Genoa, Italy

## Abstract

Several influential works propose that the acquisition of motor behavior involves different learning mechanisms in the brain, in particular supervised and reinforcement learning, that are respectively associated with cerebellar-thalamocortical and basal ganglia-thalamocortical networks. Despite increasing evidence suggesting anatomical and functional interactions between these circuits, the learning processes operating within them are studied in isolation, neglecting their strong interdependence. This article proposes a bio-inspired neurorobotic model implementing a possible cooperation mechanism between supervised and reinforcement learning. The model, validated with empirical data from healthy participants and patients with cerebellar ataxia, shows how the integration of the two learning processes could lead to benefit both learning performance and movement accuracy.

## Introduction

Supervised and reinforcement learning are two critical brain mechanisms that depend on different driving forces [1–4]. In supervised learning (SL), an agent, for example implemented with a neural network, produces an output to respond to an input pattern, for example it produces an action to respond to a visual pattern. Following this output, an external agent or a suitable internal component of the learning agent, defines a desired output pattern that represents the ‘teaching signal’ guiding the learning process, for example the correct action that was supposed to be associated with the visual pattern. The discrepancy between the agent’s output and the teaching signal defines the ‘error signal’ that is used by the agent to update its parameters, for example the connection weights of the neural network, to learn how to produce the desired output in correspondence to the input pattern [5–7].

In reinforcement learning (RL), the effect produced by the agent’s action is evaluated based on a reward obtained from the environment or autonomously computed by the agent based on the evaluation of the outcome of its action. In the early phases of learning, the agent randomly explores the environment to obtain a reward. The discrepancy between the predicted future rewards and their actual experienced values – the reward prediction error – drives the learning process [8, 9]. In particular, the agent, for example based on a neural network, uses such error to modify its connection weights in order to produce with a higher probability the actions that lead to a reward and to improve its prediction of the reward itself. Unlike SL, in RL the feedback given to the agent is less informative. For example, while learning to ride a bicycle, a SL agent could be directly supplied the correct movements to keep the bike balanced. Instead, a RL would receive a reward indicating only the global success of its actions.

The cerebellar-thalamocortical and the basal ganglia-thalamocortical loops represent the main neuronal networks underpinning, respectively, SL and RL mechanisms [1, 3, 10, 11]. In computational terms, a supervised feed-forward neural network could capture the essence of the SL mechanisms allowing the cerebellar-thalamocortical circuit to predict sensory-motor consequences of movements based on a given motor command [6, 12–14]. Instead, an actor-critic model could mimic the RL processes guided by dopaminergic feedbacks involving the basal ganglia-thalamocortical network [15–19]. In this model, the actor component is responsible for selecting motor actions depending on current sensory inputs, while the critic component produces the step-by-step signals based on the final reward, guiding the learning processes via dopamine signals [9, 16, 20–22].

In the last decades, increasing evidence has shown the existence of anatomical and functional interactions between these circuits, supporting the hypothesis that they work as a fully integrated system [4, 7, 23–25]. For example, research has led to the discovery of reciprocal synaptic projections between basal ganglia and cerebellum [26, 27] and dopaminergic projections from the ventral tegmental area and substantia nigra pars compacta to the cerebellar cortex [28–30]. Despite these evidence, the learning processes operating within these circuits are still studied in isolation, neglecting their possible interdependence.

This article proposes a computational model that, for the first time, implements a possible cooperation mechanism between the learning processes operating within the cerebellar-thalamocortical and basal ganglia-thalamocortical loops. The model is validated with a simulated bio-mimetic arm and task addressing kinematic data on motor behavior collected with healthy participants and patients affected by cerebellar ataxia [31]. The results show how the integration of SL and RL could lead to benefit both learning performance and accuracy.

## Materials and methods

### Model overview and biological underpinnings

Fig.1 shows the main elements of the model formed by a bio-constrained 2D arm model moving on the plane to reach some targets, and a neural architecture inspired by the anatomy of the brain underlying motor movement acquisition and expression. The model is formed by two macro-components. A fist component is an actor-critic RL model [19, 32], corresponding to basal-ganglia. In turn, this component is formed by an ‘actor’, that progressively develops movements by trial-and-error, and a ‘critic’, that guides such learning (and also own learning) through a reward-based signal (*δ*) elaborated at each step and corresponding to dopamine. A second component, corresponding to the cerebellum, is a feed-forward perceptron that learns movements with a SL process based on the movement errors computed with respect to the movements expressed by the whole system (actor-critic and perceptron controlling the movement in concert). The main idea underlying the model is that in early stages of movement learning, RL plays a key role to find gross solutions to the task and to favour SL, while in later stages SL plays an increasingly important role to refine moments and also to guide RL.

**Fig 1.**
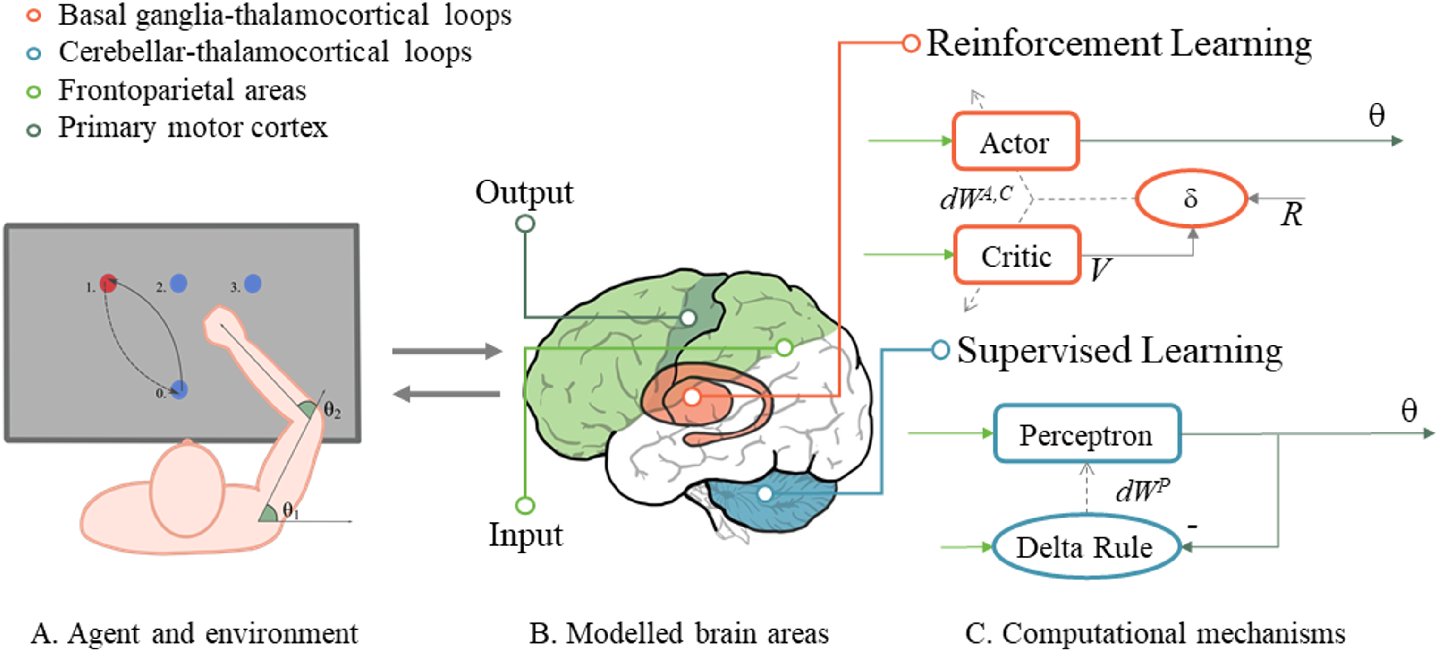
Model simulated agent and correspondences of its macro components with the brain system-level anatomy. Panel A: the simulated agent explores the environment and learns to reach the targets that activate one after the other. Panel B: brain cortical-subcortical loops that are mainly involved in the performance and learning processes underlying the behaviour. Panel C: it is possible to link the computational processes of the brain components belonging to the loops with the specific elements of the integrated model. *dW^P^*: Perceptron weights change; *dW^A,C^*: actor and critic weights change.

Substantial empirical evidence supports this idea, in particular showing that basal-ganglia and cerebellum work in a close integrated fashion to support the acquisition and expression of motor behaviour. Empirical evidence shows an early activation of the striatal nuclei during learning, followed by a later activation of cerebellar regions [33–36]. In addition, empirical evidence and theory suggest that dopaminergic signals encoding rewards could shape the coarse-grained neural patterns underlying targeted movements expressed by the basal ganglia, while at the same time furnishing a learning signal to the cerebellum [37, 38]. The teaching signal used by the SL mechanism operating in the cerebellum might thus be given by the combination of the motor noise signal from basal ganglia [39] and the dopamine signal linked to the goal of the action [37]. The first signal might be conveyed by the recently discovered subthalamic-pontine-cerebellar connection [26] and might modulate the cerebellum sensitivity to incoming error signals [4, 24]. Regarding the second signal, recent evidence shows that cerebellar granule cells encode expected reward [40] and that reward and punishment differentially influence motor learning [41, 42]. Finally, evidence supports the model hypothesise that SL processes taking place in the cerebellum retroactively inform the RL processes involving basal ganglia. This idea is supported by the discovery of a disynaptic connection linking cerebellum to the striatum via the thalamus, the cerebello-thalamo-striatal pathway [34, 43, 44].

The model is validated with data on healthy participants and patients with cerebellar ataxia. Cerebellar ataxia arises from different genetic or acquired etiologies that ultimately lead to a cerebellar impairment. Patients with this dysfunction suffer from different motor disorders such as limb incoordination and gait instability [45]. The model is in particular used to investigate how the alteration of the interplay between RL and SL computational processes might underlie the expression of the movement features observed in cerebellar ataxia. The system-level approach used here to investigate these processes agrees with recent studies supporting the importance of considering the interaction between cerebellar-thalamocortical and basal ganglia-thalamocortical circuits to understand the neural processes behind cerebellar ataxia and to discover new treatments for it [46].

### Simulated task

The model was tested with a simulated task reproducing the experimental protocol used in [31] to study cerebellar ataxia. This task consists of performing a series of reaching movements toward three circular targets (Target 1, 2, 3), turned on one after the other, starting from a resting position (Target 0). All the targets (1 cm size) are placed in a row every 17 cm (Fig.2). Each movement has to be carried out as fast and as straight as possible back and forth.

**Fig 2.**
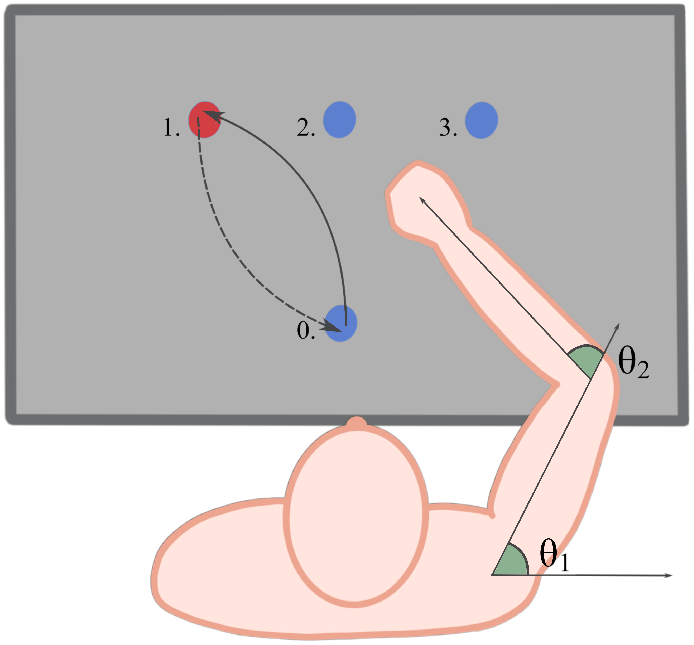
Experimental set-up and task. A simulated bio-mimetic human arm learns to reach three targets (Target 1, 2, 3) placed on a table in front of it starting from a home position (Target 0). The arm is formed by two links and is controlled in terms of the angles *θ* 1, θ2.

The model simulation was formed by 5,040 epochs. Each epoch was formed by seven trials, each consisting of 150 simulation steps. At the beginning of each epoch, the simulated upper limb rested on the starting rest position shown in Fig.2. During the first trial the model learned to move towards the turned-on home target closer to the agent (Target 0) and then during the following six trials it learned to move towards the other illuminated targets, and back to the home target, in sequence. In particular, the targets were turned on in all epochs following the fixed sequence [0,1, 0, 3, 0, 2,0]. The first 5,000 epochs formed a training phase during which the end-effector of the model learned to reach all the targets placed in the scene using only RL (henceforth called ‘RL model’) or combining the SL and RL processes (‘integrated model’). This learning phase allowed the model to learn all reaching movements performed to hit all targets. The test phase, composed of the last 40 epochs, was used to extract a series of quantitative indexes as those commonly used to describe human movement coordination as done in [31].

For each trial of the test, the indexes were computed as follow:

- *The linearity index*, measuring the trajectory curvature, was determined as the percent of increment of the total lenght of the end-effector path with respect to the minimal linear trajectory needed to reach the target:

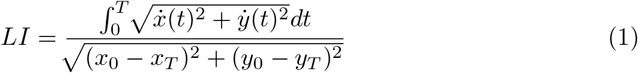
- The *asymmetry index*, related to the shape of the speed profile of the end-effector linear velocity, was computed as the ratio of the deceleration phase duration *t_post_* on the acceleration phase duration *t_pre_*:

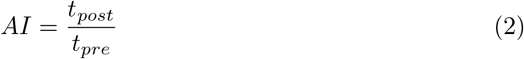
- *The smoothness index*, related to the movement jerk (third time derivative of hand trajectory), was calculated as the logarithm of the squared norm of the jerk normalized on time and space [47]:

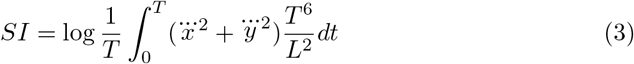

We collected data from 10 different simulated participants. The different participants were obtained by running the model with different seeds of the random number generator which in turn caused different values of the initial connection weights of the model neural networks and the exploration noise.

### Human bio-mimetic arm

The model controlled a simulated dynamical arm [47] implemented with the realistic bio-mechanical parameters as proposed in previous works [19, 48]. The shoulder position was anchored to the [0.0, 0.0] coordinates in the task space. The model could move the simulated arm changing the shoulder (*θ*_1_) and the elbow (*θ*_2_) joint angles. At the beginning of each epoch the arm had a default starting configuration (*θ*_1_ = 45.0°, **θ*_2_* = 90.0°). The range of motion of the two articular joints was [−60.0°,150.0°] (with respect to the line parallel to the table) for the shoulder, and [0.0°,180.0°] (with respect to the line formed by the upper arm) for the elbow. The time step (△t) used to numerically integrate the dynamical equations of the arm model was set to 0.01*s* and the integration was computed using a Runge-Kutta fourth order method. Finally, the muscular visco-elastic properties of the human arm were reproduced on the basis of a *proportional derivative controller* (PD controller) [19]. This mathematical model computes the torques vector (A) starting from the current angular position (*θ* = (*θ*_1_, *θ*_2_)), the desired angles (*θ_d_*) supplied by the artificial neural networks, and the current angular velocity 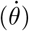:

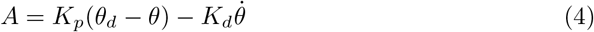

where *K_p_* was the proportional parameter (shoulder = 20.0*Nm/rad*; elbow = 10*Nm/rad*) and *K_d_* was the damping parameter (shoulder = 1.5*Nm/rad*; elbow = 1.0*Nm/rad*). These value were chosen according to human arm visco-elastic parameters [49].

### Neural architecture: functioning and learning

The architecture of the model reproduced the main brain networks and processes underlying motor learning [3, 7, 13, 19, 50], in particular the RL processes associated with basal ganglia-thalamocortical circuits and the SL processes associated with cerebellar-thalamocortical loops (Fig.1, panel B). The RL processes were implemented with an actor-critic model whereas the SL processes were implemented with a single-layer perceptron (Fig.1, panel C). These two components acted in concert to learn to produce the movements directed to reach the targets (Fig.1, panel A).

In the experiments, we compared the learning performance of the model trained only on the basis of the RL processes (the ‘RL model’, not involving the cerebellar function) with the performance of the integrated model combining both SL and RL processes (the ‘integrated model’). When we used the RL model, the output signal controlling the arm was produced by the actor component, whereas when we used the integrated model the output signal controlling the arm was a combination of the output of both the actor and the perceptron. In both cases, the output layer was composed of two units that encoded the two arm angle joints. For each target, the model trained a different neural RL and SL module in order to simulate the different population codes observed in primates’ brain motor systems when engaged in performing movements directed to different targets [51–53].

The input to the system was encoded in two different bi-dimensional neural maps reacting respectively to two scaled variables corresponding to the x-y position of the visual target (21×21 units; Fig.3), and to two scaled variables corresponding to the two proprioception angles of the arm joints (31×31 units; Fig.3). This input simulated the functioning of the thalamocortical receptive fields associated with the visual and somatosensory cortices and resulting from unsupervised learning processes (Fig.3). According to the ‘population codes’ approach, that mimics the activation of such cortical areas [54–56], each unit of each map was assumed to be located on a vertex of a 2D square grid overlapped to the neural map, where the map was assumed to encode the two variables with its two dimensions. Each unit maximally activated in correspondence to the two input values, related to the two variables to be encoded, corresponding to its vertex. This activation was lower if the two input values were different with respect to the unit preferred values, in particular it decreased progressively as a Gaussian function of their Euclidean difference (the Gaussian function had a height of 1 and a standard deviation 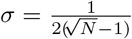, where *N* was the number of units in the map). The units of the two maps were then encoded in a whole vector to simplify the feeding of their activation pattern into the downstream model neural networks, thus henceforth we will often refer to the overall input as the ‘input layer’.

**Fig 3.**
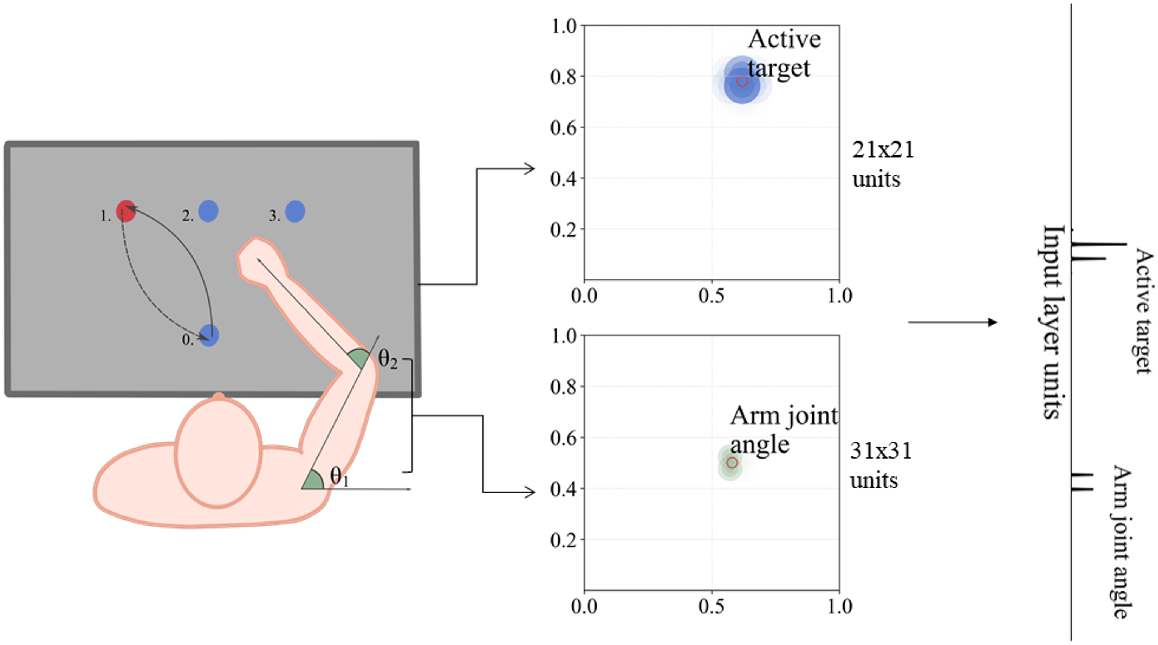
The neural encoding of the visual and proprioceptive input information. The RL model and the integrated model received two types of input data. The first one was associated with vision, represented by the coordinates of the target to reach. The second one was associated with the arm proprioception corresponding to the shoulder and elbow joint angles. These two signals were enrolled and concatenated to form the model ‘input layer’: the picture on the right shows an example of activation of this layer.

### Basal ganglia: the RL model

The RL component was formed by two sub-components, the actor and critic. These received the activation of the input layer encoding the target position and the arm proprioception (Fig.4). The actor was composed of two output units each formed by the leaky integrator model usable to mimic the activation of neural populations [57, 58]. This model had the proper level of abstraction to study system-level phenomena driving the acquisition of motor reaching skills [59]. The units of the input layer projected to the actor output units through all-to-all connections. The activation potential 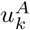 and the activation 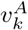 of each one of the two output units of the actor were computed as follows:

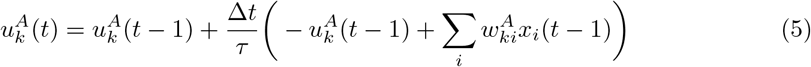

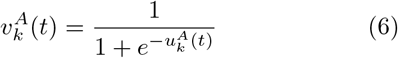

where, *x_i_* is the activation of the input unit *i*, 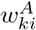 is the connection weight between input unit *i* and output unit *k*, Δ*t* (set to 0.01) is the time step of the simulation used in the Euler approximation of the model equations, *τ* (set to 0.15) is the time constant tuning the speed of the unit dynamics, and 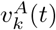 was the unit activation computed with a logistic function squashing the output in the range [0.0,1.0]. The initial connection weights were set to random values drawn from a normal distribution (*μ* = 0.0, *σ* = 0.01). The high value of *τ* was chosen so that the units’ dynamics reflects the information processing by the basal ganglia that is faster then the cerebellar one [60, 61].

**Fig 4.**
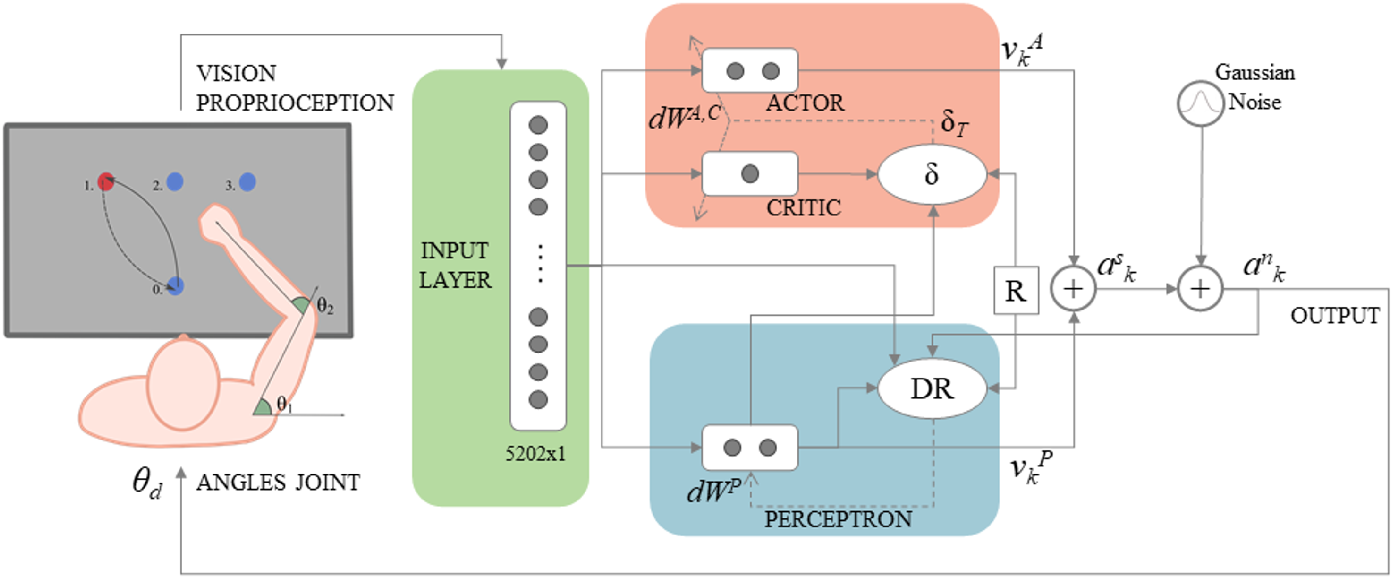
A schematic view of the integrated model components interaction. Both model components, the RL component (red box) and the SL component (blue box), received the input composed of the visual and proprioceptive information and controlled the bio-mimetic arm movement. DR: Delta Rule; *dW^P^*: perceptron weights change; *dW^A,C^*: actor and critic weight change.

A noise value *n_k_*(*t*) was added to each model output unit to support the exploration requested by RL: 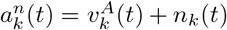. The noise value *n_k_*(*t*) depended on a first order filter, which avoided that the effect of noise was cancelled by the inertia of the dynamic arm [19], and on the success rate of the model to reach each target, which ensured a more stable performance with the progression of learning [62]. The noise was in particular computed as follows:

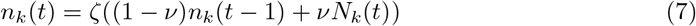

where *N_k_*(*t*) was a noise value extracted from a normal function with zero mean and a standard deviation set to 0.5, and *ν* (*ν* = 0.198) was the time constant of the first order filter; the parameter *ζ*, ranging in [0.0, 1.0], was computed as a moving average of the success of the model for the given target in the last 600 trials, where the success was encoded with a binary value in [0.0,1.0].

The critic component had one output unit with which it evaluated the outcome of the action selected by the actor. As for the actor, the input layer projected to the critic output unit through all-to-all connections initially set to random values drawn from a normal distribution (*μ* = 0.0, *σ* = 0.01). The output unit computed the activation potential as for the actor units but used a linear transfer function, rather than a logistic one, to compute its activation.

The connection weights of the critic were updated as illustrated below so the output unit activation *V*(*t*) encoded an estimate (‘evaluation’) of the future reward achievable within the trial from the current state. The evaluation was in particular defined as the expected discounted future reward *γ*^0^*r*(*t* + 1) + *γ*^1^*r*(*t* + 2) + *γ*^2^*r*(*t* + 3)… where *γ* was the discount factor (0 < *γ* < 1, set to 0.99). At each simulation step, if the end-effector was on the target, the model received a reward. The reward *R*, ranging in [0.0, 1.0], was equal to 0.0 when the end effector was off the target (target now yet achieved), and in case of success it was computed with an inverse relation with respect to the impact speed with the target:

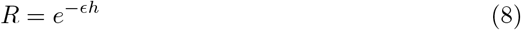

where *ϵ* (set to 0.2) was a parameter and *h* was the end-effector linear speed (*m/s*). According to this formula the reward was higher for a lower contact speed close to zero, and decayed with the rate *ϵ* with a higher contact speed. This reward favoured the learning of a slowly contact with the target as it happens in humans [19].

The reward *R*(*t*), and two succeeding evaluations *V*(*t*) and *V*(*t* – 1), were used to compute the *temporal difference error*, *δ*(*t*), used in each step t to train both the actor and the critic [9]. The computation of *δ*(*t*) was based on the episodic nature of the RL process considered in the target experiment:

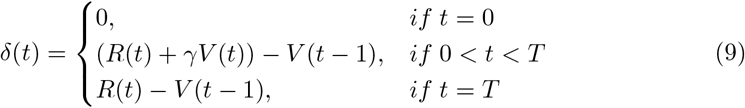

Based on this formula, in the first step of each trial (*t* = 0) *δ*(*t*) was set to 0.0 as there was not a ‘previous evaluation’ to guide training, so training did not take place. On the other hand, at the end of the trial (*t* = *T*) the Δ_*t*_ was equal to the difference between the final reward, *R*(*t*), and the previous evaluation, *V*(*t* – 1), while *V*(*t*) was set to zero (because the trial ended and so there were no ‘future rewards’). In all other steps, the formula used the standard temporal difference learning rule.

The learning signal *δ*(*t*) was used to train both the critic and the actor. The critic connection weights 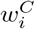 were in particular updated as follows:

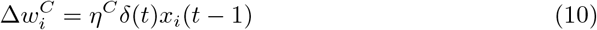

where *η^C^* (set to 0.06) was the critic learning rate. This formula led the critic to improve its evaluation *V*(*t*) of future rewards. The actor connection weights 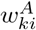 were instead updated as follows:

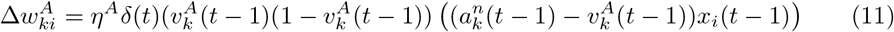

where *η^A^* (set to 0.6) was the actor learning rate, and 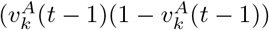 the derivative of the output unit logistic function. This formula led the actor to increase the probability of producing the output 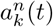 in case of positive *δ*(*t*), and to decrease it otherwise.

### Cerebellum: the SL component

In line with literature supporting the fast signal processing of the cerebellar-thalamocortical circuit with respect to the basal ganglia-thalamocortical circuit [63–67], the perceptron differed from the actor-critic component for the time integration parameters used. In particular, the perceptron computed the output signal, 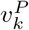, according to a leaky integrator model and a logistic function similar to the one used for the actor (Eq. 5). However, it differed from the actor for the time parameter *τ^P^* that was here set to 0.01 (Δ*t* was set to 0.01 as for the other components of the model). This value of *τ^P^* was chosen to make the SL component capable of processing signals at a faster temporal scale facilitating the fine-tuning of motor movements [60, 61].

During the trajectory execution, the memory trace about the value 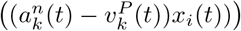 involving the input state *x_i_*(*t*), the associated perceptron output 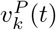, and the model output 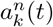, was stored in a memory. In the brain, this memory could correspond to a dynamic memory or a chemical trace. At the end of the trial, the perceptron connections were updated as follows based on the memory trace and the reward *R*:

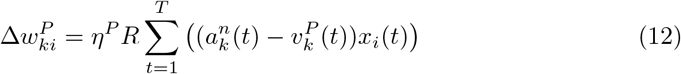

where *η^P^* (set to 0.6) was the perceptron learning rate. In this way, from the first time in which the model reached the target under the guidance of the RL process, the produced sub-optimal motor trajectory 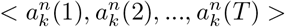 furnished the teaching signals to train the perceptron.

### The integrated model

Each output unit of the integrated model, 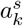, was activated as the arithmetic average of the activation of the actor output unit 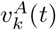 and of the perceptron output unit 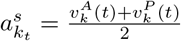. The output was also added a noise, analogously to what done in the RL model, to compute the overall integrated model noisy output as follows: 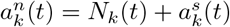. The contribution to the overall output of both the actor and the perceptron was a crucial difference with respect to the RL model. At the beginning of the simulation, the SL component did not produce any benefit to the exploration strategy. At this stage, the weights of both the actor and the perceptron components were set to random values and so they were not able to reach the targets. As soon as the model got rewarded the first time, the SL component acquired the (initially coarse) trajectory leading the arm end-effector to reach the target. On the other hand, the influence of the SL component on the exploratory behaviour supported the RL module training. This because since the first trial in which the model received the reward, the output signal of the SL component supplied an initially gross, but gradually improving, trajectory to reach the target, and this affected the noisy exploration of the model making it more focused around the area surrounding the target. The actor training equation of the integrated model was modified to take into account this influence of the SL component over the RL one as follows:

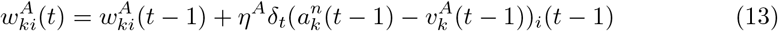

To simulate ataxia, we damaged the SL output by adding noise to it. In particular, at each simulation step the *τ^P^* value of the perceptron was drawn from a normal distribution (*μ* = 0.15, *σ* = 0.002; notice that in this way the cerebellum becomes as slow as the basal-ganglia actor having a *τ^A^* = 0.15). A further noise drawn from a normal distribution (*μ* = 0.0, *σ* = 0.03) was added to the perceptron output. This impairment affected the functioning and training processes of both the RL and SL modules.

## Results and Discussion

This section shows and discusses the results obtained with the RL model and the integrated model in order to: (a) compare the learning performance of the two models and compare their capacity to reproduce the data from human healthy participants; (b) validate the capacity of the integrated model to reproduce the differences between patients with cerebellar ataxia and healthy participants.

### Learning performance

The integrated RL-SL model exhibits a faster learning process with respect to the RL model (Fig.5). This faster learning process of the integrated model was due on the improved exploration strategy it followed. In particular, since the model gets the reward for the first time, the SL component can learn a first coarse trajectory to reach the target. This trajectory affects the exploration reducing its randomness and makes it more focused on the target (see small circles in Fig.5). The RL component then gradually improves the sub-optimal initial solution trial after trial while avoiding unneeded explorations as instead it happens in the pure RL model. Similarly, the SL component gradually improves its behaviour because it receives a better teaching signal (trajectories) from the RL processes. This mutual training between SL and RL thus produced a faster learning process.

**Fig 5.**
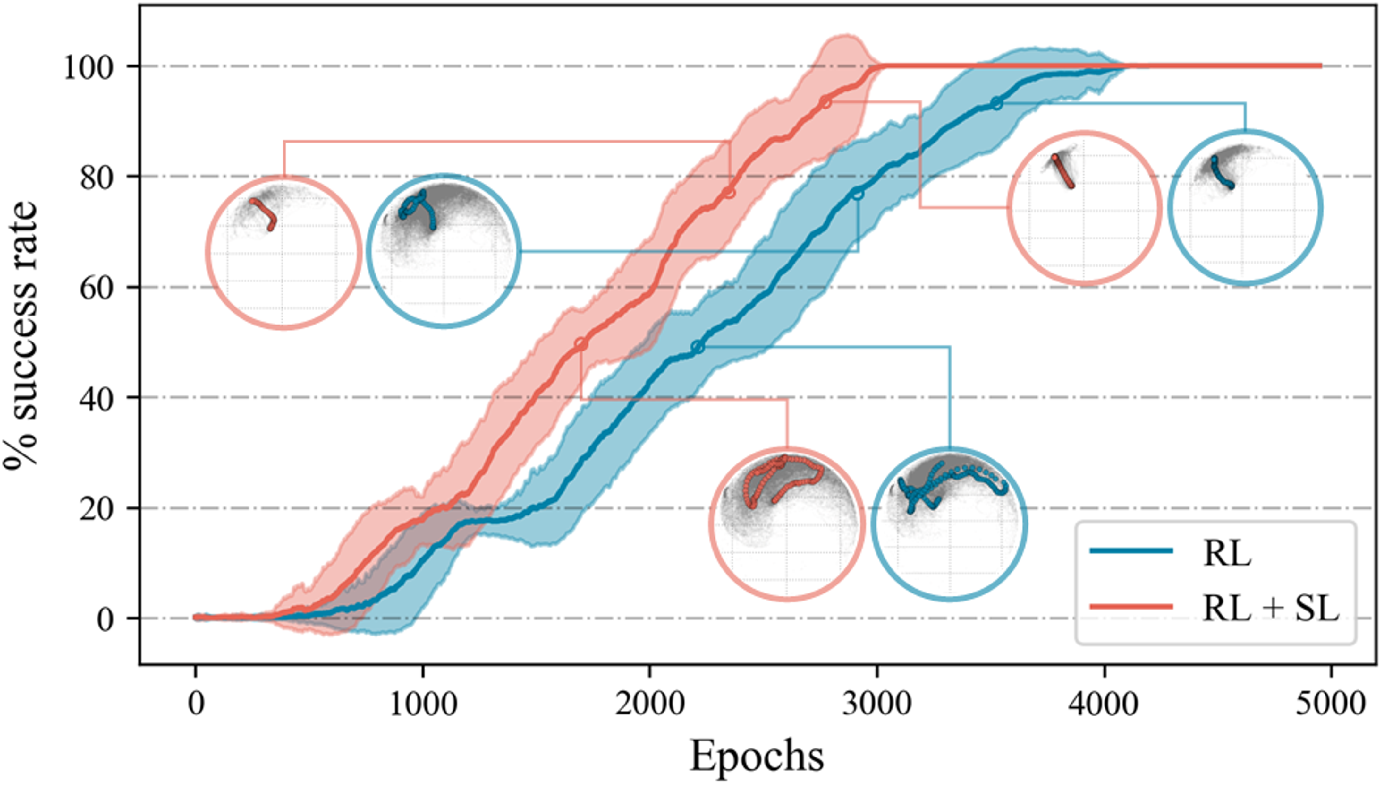
Learning performance of both the integrated RL+SL model and the RL model. The circles represent the area (grey cloud) explored during the learning epochs when the models reached a success rate of 50%, 80%, and 100%. Each circle also shows examples of trajectories actually followed by the end effector during the movement. Notice the larger exploration are of the RL model indicating a more erratic behaviour with respect to the integrated model.

The test indicates that the more focused exploration strategy supported by the SL component was critical to speed up the overall learning process. Based on this result, we can speculate that a better learning performance could be one of the potential advantages which led to the evolutionary emergence of the integrated cerebellar-thalamo-cortical and the basal ganglia-thalamo-cortical system.

### Addressing data of healthy participants

Fig.6 compares the ability of the RL and RL+SL models to reproduce data on healthy participants’ motion indexes [31]. The figure shows that the two models have a statistically significant difference in terms of linearity and smoothness but do not differ in terms of asymmetry. For each index and for each model we also computed a ‘discrepancy index’ (Δ(*SIM, REAL*)) representing the absolute value of the difference between the simulated data and data from real healthy participants studied in [31]. The values of the index are reported in Table 1. The Table shows that the discrepacy is smaller for all the values of the indexes of the integrated RL+SL model with respect to those of the RL model. These results show that the integrated model produces a behaviour with features that are closer to those of the behaviour exhibited by the real participants of the experiment with respect to the single-mechanism RL model.

**Fig 6.**
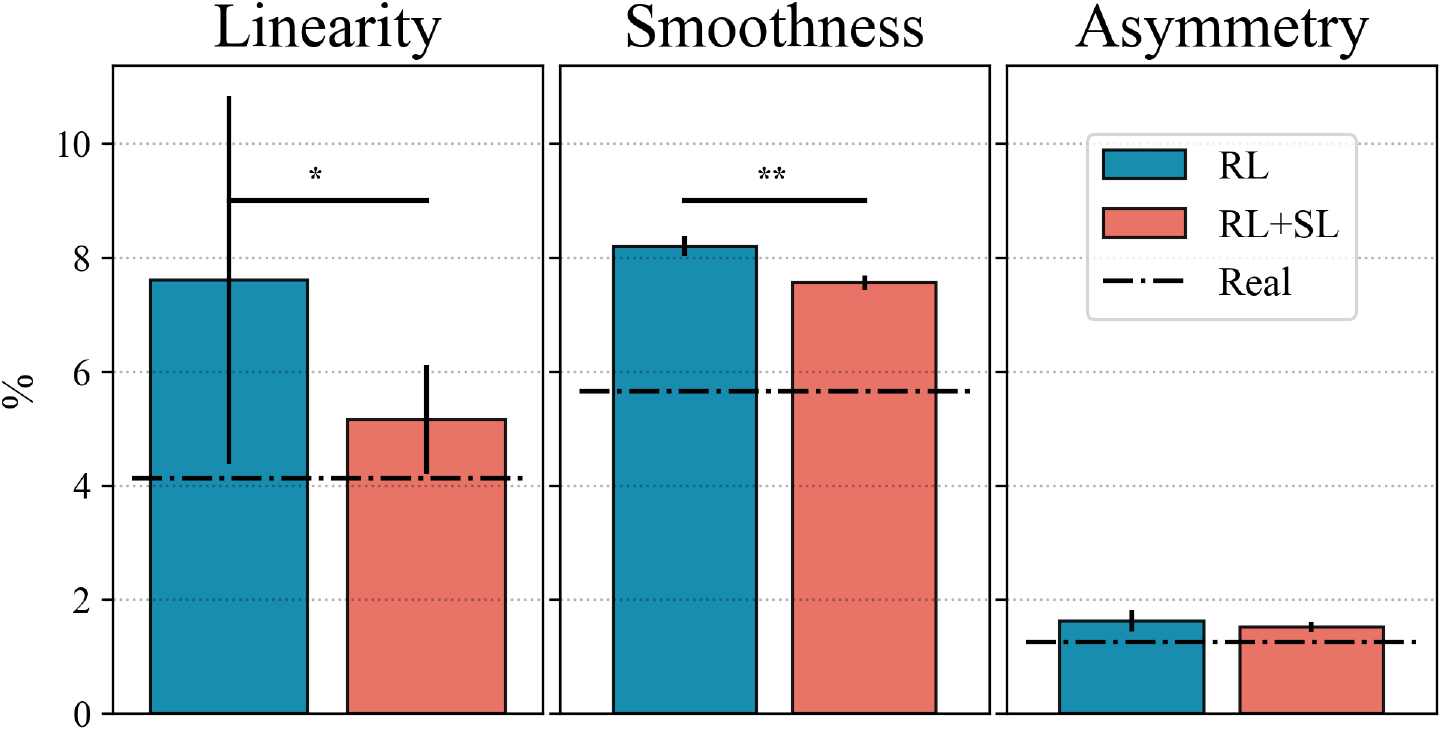
Motion indexes of the RL and RL+SL models. Both linearity and smoothness indexes showed a statistically significant difference between the two models (for the two indexes were respectively 0.042 (* p-value < 0.05) and 7.28 ·10^-8^ (** p-value < 0.001)). The black dotted lines represent the values of the indexes of the real healthy participants.

**Table 1.** Statistical analysis comparing the behavioural indexes produced by the two RL+SL and RL models, and the indexes produced by the models and those produced by real participants. The p-values refer to the t-tests and show that there is a statistically significant difference between the linearity and smoothness indexes reproduced by the two models (highlighted in bold) while the two models do not statistically differ in terms of asymmetry. The discrepancy index (Δ(*SIM, REAL*)) indicates the absolute difference between the mean values of the indexes of the models reported in the table and those of the real participants (the smaller value between the two models is highlighted in bold).

| Motion index | Mean ± Std | p-value | $\Delta(SIM, REAL)$ |
| --- | --- | --- | --- |
| Linearity RL | $7.61 \pm 3.4$ | **0.042** | 3.48 |
| RL+SL | $5.16 \pm 1.0$ | | **1.03** |
| Smoothness RL | $8.2 \pm 0.19$ | **$7.28 \cdot 10^{-8}$** | 2.54 |
| RL+SL | $7.56 \pm 0.13$ | | **1.90** |
| Asymmetry RL | $1.63 \pm 0.2$ | 0.128 | 0.38 |
| RL+SL | $1.52 \pm 0.1$ | | **0.27** |

### Addressing data on participants with cerebellar ataxia

Fig.7 shows a comparison between the values of motion indexes of the intact RL+SL model simulating the healthy participants, and the values of the same indexes scored by the model where the SL component was lesioned after learning to simulate the patients with cerebellar ataxia. The figure shows that the lesioned model exhibits movements that are less linear, less smooth, and more asymmetric with respect to the intact model. These results reproduce the qualitatively difference that can be seen between real patients with cerebellar ataxia compared to human healthy participants [31].

**Fig 7.**
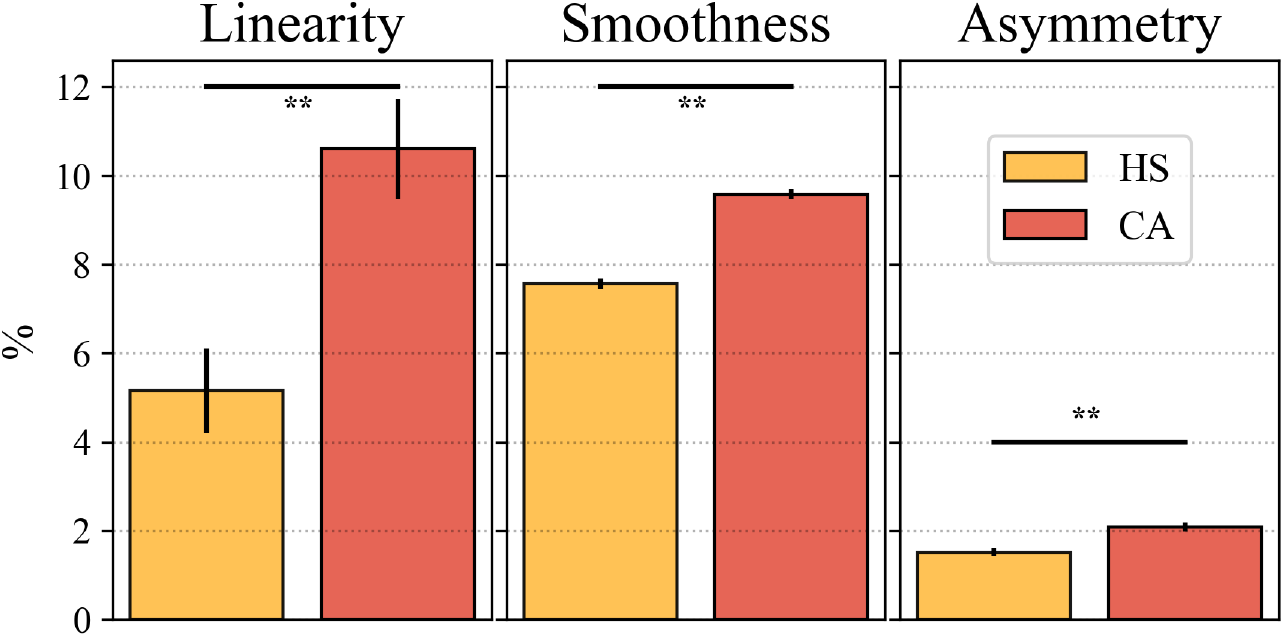
Simulation of healthy participants (HS) and cerebellar ataxia patients (CA) with the integrated RL+SL model. The t-tests between the values of the indexes show a statistically significant difference between the two model conditions (the p-values for the thee indexes were respectively: 2.43 ·10^-9^, 5.27 ·10^-10^, and 5.36 ·10^-18^).

The relevance of these results resides in the fact that computational approaches proposed in the literature to address human motor acquisition mainly rely on one learning process [19, 32, 68, 69]. However, this single-mechanism approach does not reflect the empirical evidence on how the brain actually learns and expresses movements, and thus could fail to model and understand relevant features of movement learning, such as its overall dynamics and specific features. Moreover, this approach does not allow the use of the same model to address different datasets involving experiments on different learning processes and brain motor components (e.g., relating to basal-ganglia, cortex, and cerebellum).

## Conclusion

The integrated model proposed in this article offers a system-level framework supporting our understanding of the brain mechanisms underlying motor learning. The key idea underpinning the model is that the brain integrates reinforcement and supervised learning processes to increase the speed of motor movements learning while at the same time ensuring a high regularity and effectiveness of action.

This idea is also relevant to technology applications, particularly since the presented model was validated with a simulated robotic plant that, although simple, had dynamics with realistic characteristics. In this respect, a relevant limitation of autonomous reinforcement learning robotic systems is its slow progression. The model and results proposed here prompt future explorations of solutions integrating SL and RL to overcome such limitation.

In terms of the study of the brain and the acquisition of motor skills, the proposed model and approach encourages an integrated interpretation of existing data proposing that human movement acquisition might rely on a mixture of reinforcement and supervised learning processes from a system-level perspective [4, 46]. In particular, the proposed mechanisms are shown to speed up movement learning and to improve movement regularity thus reproducing the features of human data related to healthy participants. Moreover, a lesioned version of the model mimicking the impairment of patients with cerebellar ataxia allowed the reproduction of the reduced performance exhibited by these type of patients with respect to healthy participants.

The current model represents a first important step to study the possible integration of reinforcement and supervised learning processes in the brain. Future versions of the model could increase the realism and the quantitative matching of data of the model by including features that reproduce the interaction between cortical-subcortical learning mechanisms [70, 71], and an architecture more closely reproducing the brain hierarchy underlying the acquisition and expression of motor movements with possibly a more explicit representation of targets [72–74].

## Acknowledgments

This research was supported by the ‘Advanced School in Artificial Intelligence’ (www.as-ai.org) and by ‘AI2Life s.r.l.’ (www.ai2life.com). This research has also received funding from the European Union’s Horizon 2020 research and innovation program under grant agreement no. 713010, Project ‘GOAL-Robots - Goal-based open-ended autonomous learning robots’.

## Authors’ contributions

**Adriano Capirchio**: Formal Analysis, Investigation, Funding acquisition, Methodology, Software, Validation, Visualization, Writing - original draft; **Chiara Ponte**: Data curation, Formal Analysis, Investigation, Resources, Software, Validation, Visualization, Writing - review & editing; **Francesco Mannella**: Data curation, Resources, Software; **Gianluca Baldassarre**: Conceptualisation, Funding acquisition, Investigation, Methodology, Supervision, Validation, Writing - review & editing; **Elisa Pelosin**: Methodology, Validation, Writing - review & editing; **Daniele Caligiore**: Conceptualization, Funding acquisition, Investigation, Methodology, Project administration, Supervision, Software, Validation, Visualization, Writing - original draft, Writing - review & editing.

## Supporting information

**S1 Fig. Model simulated agent and correspondences of its macro components with the brain system-level anatomy.**

**S2 Fig. Experimental set-up and task.**

**S3 Fig. The neural encoding of the visual and proprioceptive input information.**

**S4 Fig. A schematic view of the integrated model components interaction.**

**S5 Fig. Learning performance of both the integrated RL+SL model and the RL model.**

**S6 Fig. Motion indexes of the RL and RL+SL models.**

**S7 Fig. Simulation of healthy participants (HS) and cerebellar ataxia patients (CA) with the integrated RL+SL model.**

